# No-take zone effectiveness in linear coastal systems depends on spatially heterogeneous placement and larval dispersal directionality

**DOI:** 10.64898/2026.02.25.707921

**Authors:** Marie Saitou, Louise Chavarie, Thrond Oddvar Haugen

## Abstract

Marine protected areas (MPAs) are widely implemented to protect and/or rebuild exploited populations, yet their population-level effectiveness remains highly variable. Key unresolved knowledge gaps include how the size and the spatial placement of no-take zones interacts with larval dispersal directionality, particularly in linear coastal systems where connectivity is asymmetric. For sedentary species with planktonic larvae, such as European lobster, it is unclear under which dispersal regimes spatial configuration of protection critically determines positive demographic outcomes. Here, we address this gap using a spatially explicit individual-based model parameterized for the European lobster. We ask (i) whether no-take zones consistently enhance abundance and size structure relative to fished areas, and (ii) whether the positioning of no-take and open areas affects spatial protection while holding total protected area constant and (iii) how the alignment between larval dispersal direction and the positioning of no-take areas influences protection outcome. We contrast local, symmetric long-distance, and strongly unidirectional larval dispersal across alternative MPA layouts with equal total protected area but with different spacing. We show that no-take zones reliably increase abundance and the prevalence of large individuals. However, when larval dispersal is strongly unidirectional, population recovery depends on reserve placement: downstream no-take zones benefit from both larval import and local retention, whereas upstream reserves primarily export reproductive output and show limited local recovery. These results indicate that reserve performance cannot be evaluated independently of connectivity structure and identify dispersal directionality as a key determinant of when and where spatial configuration matters for MPA effectiveness in linear coastal systems.

## Introduction

Marine protected areas (MPAs hereafter), and particularly no-take zones, are widely used tools in fisheries management to mitigate the impacts of harvesting and rebuild depleted populations (Sala and Giakoumi, 2018; Coppa *et al*., 2021; Erm *et al*., 2024). While numerous empirical studies report increases in local abundance and body size within protected areas, the magnitude and spatial extent of these benefits vary considerably among systems (Stobart *et al*., 2009; Ovando *et al*., 2021) and species (Batista *et al*., 2015; Sullivan-Stack *et al*., 2024). Such variability is often attributed to species-specific life-history traits, spatial habitat structure, and patterns of connectivity driven by larval dispersal (Krueck *et al*., 2017; Cristiani *et al*., 2024). For sedentary coastal species with planktonic larvae, such as the European lobster (*Homarus gammarus*), protection of adult individuals alone may be insufficient to generate population-level benefits beyond reserve boundaries if larval export and settlement are strongly spatially structured (Øresland and Ulmestrand, 2013). In linear coastal systems such as fjords and archipelagos, where dispersal may be directional rather than symmetric, the spatial placement of no-take zones relative to larval flow has the potential to critically shape management outcomes (Woodson, 2018). In such systems, no-take zones may function as demographic sources that produce larvae, while adjacent fished areas act as sinks where larvae settle, grow, and are subsequently harvested(Conklin *et al*., 2018; Clavel-Henry *et al*., 2024). Yet, this interaction remains not fully resolved in a mechanistic, life-history–explicit framework (Heino *et al*., 2013).

Empirical work on European lobster conservation has consistently reported positive local responses to protection, including increased density, biomass, and size structure within no-take zones (Moland *et al*., 2013a, 2013b; Fernández-Chacón *et al*., 2021; Perry *et al*., 2025). Long-term monitoring programs have shown that reduced fishing mortality allows individuals to survive to larger sizes and higher reproductive output (Moland *et al*., 2013b; Knutsen *et al*., 2022). However, these effects are often spatially restricted to protected areas and adjacent habitats, with limited evidence for consistent population-level recovery at broader spatial scales (Moffitt *et al*., 2011; Nillos Kleiven *et al*., 2019). This limitation is commonly attributed to the interaction between sedentary adult behaviour and a pelagic larval phase, where successful replenishment of fished areas depends on the magnitude and direction of larval dispersal (Hamilton *et al*., 2021). Although larval connectivity is widely recognised as a key determinant of reserve performance(Saenz-Agudelo *et al*., 2011), most existing studies either infer connectivity indirectly or focus on local responses without explicitly examining how dispersal structure interacts with the spatial arrangement of protection. Previous theoretical and applied studies have demonstrated that simplified patch-based models can be sufficient to isolate key mechanisms underlying MPA performance, particularly when the objective is to disentangle the effects of dispersal structure and reserve placement rather than to reproduce site-specific hydrodynamics (Smith and Jensen, 2008; Harrison *et al*., 2020; Barceló *et al*., 2021; Backus *et al*., 2022; Quennessen *et al*., 2023; Hopf *et al*., 2024). Studies in terrestrial landscapes have shown that protection organised as spatially heterogeneous mosaics of multiple patches can enhance population persistence and connectivity more effectively than single large reserves (Law and Dickman, 1998; Fahrig *et al*., 2011; Ricketts and Sandercock, 2016; Regolin *et al*., 2020). In contrast, whether comparable principles extend to marine no-take zones remains relatively unexplored, despite the importance of larval dispersal in connecting protected and exploited areas.

To address this gap, we use a spatially explicit individual-based model to examine how larval dispersal directionality and the placement of no-take zones jointly shape population dynamics in a sedentary coastal species. The model represents essential biological processes of the European lobster, including growth, maturation, sex-specific reproduction, density-dependent recruitment, and size- and egg-based fishing regulations (Øresland and Ulmestrand, 2013; Haugen *et al*., 2023), while deliberately simplifying spatial structure to a one-dimensional coastal system. By contrasting alternative dispersal regimes, and by comparing different configurations of protected and fished patches with equal total protected area, we isolate the mechanisms by which connectivity and spatial management interact. Specifically, we ask (i) whether no-take zones consistently enhance abundance and size structure relative to fished areas, and (ii) whether the positioning of no-take and open areas affects spatial protection while holding total protected area constant and (iii) how the alignment between larval dispersal direction and the positioning of no-take areas influences protection outcome.

## Materials and methods

### Model overview

We developed a spatially explicit individual-based model to investigate how larval connectivity, the spatial configuration of marine protected areas (MPAs), and fishing regulations influence population structure and reproductive output. The conceptual structure of the individual-based model, including the life cycle, larval dispersal regimes, spatial domain, and fishing regulations, is summarized in **Figure 1**. The model represents annual population dynamics of a sedentary coastal species along a linear coastline, with between-patch connectivity driven exclusively by planktonic larval dispersal and spatially explicit fishing management. While parameterized using traits of European lobsters (*Homarus gammarus*), the model is intended as a general framework for evaluating spatial protection strategies.

**Figure 1.**
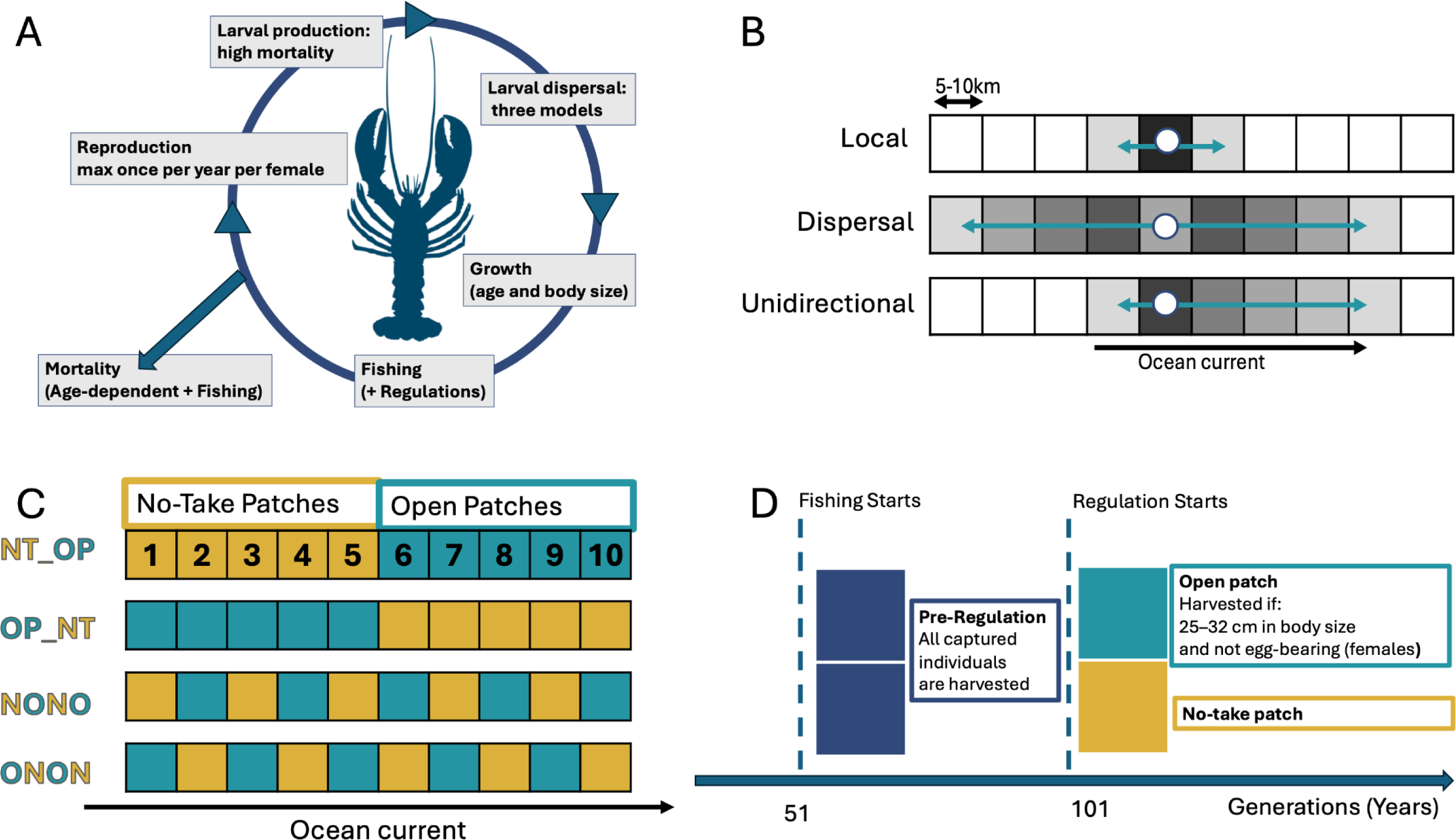
Conceptual structure of the lobster life-cycle model, dispersal scenarios, and spatial management layouts. **(A) Life-cycle structure.** Panel A illustrates the annual life cycle of the individual-based lobster model. Within each one-year time step, mature females may attempt reproduction probabilistically, producing larvae subject to high early mortality. Larvae disperse along a linear coastal system according to one of three dispersal scenarios. Surviving individuals settle, grow in age and body size, may move between adjacent internal patches as adults, and experience age-dependent natural mortality as well as fishing mortality subject to management regulations. All processes are updated annually in a fixed order. **(B) Larval dispersal scenarios.** Panel B summarizes the three larval dispersal scenarios considered in the model. These scenarios differ in dispersal distance and directional bias: a *Local* scenario with strongly localized retention, a *Dispersal* scenario with symmetric bidirectional spread over multiple patches, and a *Unidirectional* scenario with strong downstream bias combined with long-distance dispersal. These scenarios are implemented using a distance-decaying dispersal kernel with a directional bias parameter and are applied uniformly across the spatial domain. **(C) Spatial domain and MPA layouts.** Panel C shows the spatial structure of the model, consisting of ten internal patches (patch 1–10) arranged along an upstream–downstream axis. Four alternative marine protected area (MPA) layouts are examined. One patch represents ∼5–10 km of coastline, corresponding to commonly inferred larval dispersal scales for European lobster and similar coastal invertebrates(Cowen and Sponaugle, 2009; Øresland and Ulmestrand, 2013). Two layouts consist of contiguous blocks of patches: one with upstream no-take patches followed by downstream open patches, and one with the reverse configuration. Two additional layouts use alternating no-take and open patches along the coast. These layouts are labeled using compact notation: **NT_OP** (upstream no-take, downstream open), **OP_NT** (upstream open, downstream no-take), **NONO** (alternating no-take and open starting with no-take upstream), and **ONON** (alternating open and no-take starting with open upstream). Each layout contains five no-take and five open patches to be investigated. **(D) Fishing regimes and regulation phases.** Panel D illustrates the fishing regimes applied to the spatial domain. Prior to the onset of regulation, all patches operate under an open-access regime with unrestricted harvest. After regulation begins, no-take patches are fully closed to fishing, whereas open patches allow harvest only within a defined size slot (25–32 cm) and exclude egg-bearing females from harvest. Fishing is implemented as a trap-based process and applies only to adult individuals.

Although the model draws on empirical knowledge from long-term European lobster monitoring, it does not directly incorporate empirical datasets beyond the dispersal component. Empirical observations (Haugen *et al*., 2023) were used to inform key structural assumptions, including low adult movement between management areas combined with frequent movement within areas (e.g. among traps), observed reductions in catchable male abundance in open areas following exploitation, and the practical details of fishing regulations such as protection of ovigerous females and size-based harvest limits. These observations were used to guide model structure rather than to reproduce specific empirical time series. Parameter values were not chosen to reproduce specific observed time series, but rather to ensure consistent population dynamics across scenarios (e.g. avoiding unrealistically rapid extinction, population burst or runaway ageing), while retaining qualitative realism. This design prioritises mechanistic insight into the interaction between larval connectivity, spatial protection, and fishing regulation over exact numerical prediction.

#### Life-cycle structure and annual dynamics

As illustrated in **Figure 1A**, we implemented a spatially explicit, individual-based model in SLiM 3.2 (Haller and Messer, 2019) to simulate annual dynamics of a sedentary coastal species with planktonic larval dispersal and size-dependent harvest mortality. Each simulation year proceeds through a fixed sequence of processes: early-life survival, aging, somatic growth, reproduction, and fishing mortality. Survival is governed by a Gompertz function (Hernandez-Llamas and Ratkowsky, 2004) calibrated to produce realistic senescence patterns. Individual body size follows a von Bertalanffy growth curve (von Bertalanffy, 1966) with heritable growth deviation.

Observed adult movement in European lobste*r* is typically restricted to within a ∼50–200 m radius. In our model, adult lobsters do not move between spatial patches once settled. All adult movement occurs within patches (e.g., among traps), but no adult crosses patch boundaries. This assumption is based on tagging data from coastal Norway (Haugen *et al*., 2023), where inter-area movements were extremely rare and, when observed, often took multiple years to occur. In contrast, larval dispersal is modeled explicitly as the sole mechanism of between-patch connectivity. Within-patch movement (e.g., among traps) is implicitly modeled through randomized trap sampling. While lobsters exhibit complex behavior including shelter fidelity and intraspecific competition (van der Meeren, 2002; Agnalt *et al*., 2017), the model operates at the patch level. Patch occupancy is governed by demographic processes only.

European lobster females often reproduce every 1–2 years due to prolonged egg incubation (9–12 months;(Agnalt, 2008; Ellis *et al*., 2015a). To reflect this observation, we implemented a simplified reproductive blocking mechanism. In the model, successful reproduction sets an internal egg flag to 2, which decrements by 1 each year. Individuals with a nonzero egg flag are not eligible to spawn. This ensures that each female reproduces no more frequently than once every two years, in alignment with observed brooding intervals. Offspring number follows a Poisson distribution, reduced by density dependence and an additional overcrowding penalty. Offspring inherit growth parameters from parents and are assigned a natal origin based on maternal location (**Figure 1A**).

#### Larval dispersal and connectivity scenarios

To evaluate the role of spatial connectivity, we implemented three contrasting larval dispersal scenarios (**Figure 1B**): Local (short-range, symmetric), Dispersal (long-range, symmetric), and Unidirectional (strongly downstream-biased). Larval dispersal occurs via a directional, distance-decaying kernel applied along a one-dimensional linear coastline composed of 12 patches (positions 0 to 11). Larvae originating from position *i* are assigned destination *j* with probability decreasing exponentially with |i−j|, and with asymmetry controlled by a bias parameter (*larvaBias*) representing the proportion of downstream dispersal. Only internal patches (positions 1–10) are analyzed; external patches (0, 11) act as larval sources/sinks but are not recorded. These scenarios differ in dispersal distance and directional bias but share the same underlying structure: larvae disperse along the coastline according to a distance-decaying kernel with an adjustable downstream bias (Cowen and Sponaugle, 2009; Planes *et al*., 2009). These scenarios are not intended to reproduce site-specific hydrodynamics; instead, they span a gradient of connectivity structures commonly invoked in theoretical and applied MPA studies.

### Spatial layout and management configurations

The model domain includes ten internal coastal patches arranged linearly (positions 1 to 10), each assigned to either a no-take zone or an open area depending on one of four spatial layouts: upstream-blocked (NT_OP; large homogeneous patches designated as protected and fished), downstream-blocked (OP_NT, large homogeneous patches designated as fished and protected), alternating starting with no-take (NONO; heterogeneous patches of protected and open areas), and alternating starting with open (ONON; heterogeneous patches of open and protected areas). In each layout, five patches are designated as no-take zones (fully closed to fishing), and five as open areas, subject to selective harvest regulations with all scenarios maintaining the same total protected area (**Figure 1C**). Spatial configuration is fixed for each simulation replicate.

### Fishing regime and regulation phases

Fishing regulations are summarized in **Figure 1D**. Fishing is implemented as a size-selective, trap-based harvest process acting on adult individuals. Prior to the onset of regulation (generations 1–100), all patches operate under open-access conditions, where any captured individual is subject to harvest regardless of size or reproductive status. Following the introduction of spatial regulation (from generation 101 onward), no-take patches are fully closed to harvest, whereas open patches enforce a harvest slot limit: only individuals between 25 and 32 cm in body length are eligible for harvest. In addition, egg-bearing females are excluded from harvest in open areas. Capture probability is size-dependent, modeled using a logistic function with individual- and trap-level variation. Each patch is assigned a fixed number of traps per year (*nTrap*), with dynamic adjustment of effort post-regulation based on prior harvest performance to approximate a constant yield target. Traps operate by sampling a fixed number of candidate individuals per patch, applying the capture probability function to each, and executing harvest only if all regulatory conditions are satisfied. Harvested individuals are removed from the population.

### Simulation protocol and outputs

Each scenario (i.e., combination of dispersal regime and MPA layout) was simulated for 200 replicate runs, using distinct random seeds. Simulations span 200 discrete generations, with fishing initiated at generation 51 and regulation implemented at generation 101. The first 50 generations allow the population to stabilize under natural dynamics. All simulations were run using SLiM 3.2 on the Orion high-performance computing cluster at Norwegian University of Life Sciences.

Population abundance was quantified as the total number of individuals per patch, stratified by area type (no-take or open to fishing) and generation, with counts recorded every 10 generations. Size structure was described using frequency distributions of individual body sizes in 5-cm bins, stratified by sex, patch, and area type, and sampled at generations 100, 120, 140, 160, and 200. Age structure was recorded concurrently with size structure and summarized as age distributions by sex and patch to allow direct comparison of demographic patterns. Natal origin was determined for each patch and sampling time by calculating the proportion of individuals originating from each natal patch, including external patches, enabling assessment of larval connectivity and dispersal among areas. In addition, a detailed trap log was maintained to document capture and harvest decisions, including information on individual size, sex, egg status, and the reason for release or harvest. These records were compiled every 20 generations to allow auditing of compliance with harvest rules and to verify that management regulations were applied consistently throughout the simulations.

All outputs are stored in tab-delimited format, with standardized headers including replicate ID, scenario label, generation, and spatial position. Summary statistics were computed across replicates, and data were visualized using boxplots, stacked bar charts, and time series plots, as described in the Results section.

### Downstream analyses

#### Population size dynamics

Total population size was calculated by summing abundances across internal patches for each replicate, generation, dispersal regime, MPA layout, and area type. Replicate-level totals were aggregated by calculating means and standard errors across replicates. Temporal analyses distinguished three simulation phases: a pre-fishing phase, a fishing phase with unrestricted harvest, and a regulated phase with spatial closures and selective harvest.

#### Spatial variation in population abundance

Spatial heterogeneity in abundance was quantified using patch-level population sizes at a focal post-regulation generation. For each replicate, abundances were summed within each internal patch. Distributions across replicates were used to compare patch-level population sizes among dispersal regimes and MPA layouts, while distinguishing between open and no-take patches.

#### Size- and sex-structured population composition

Size-structured population composition was analyzed using size-binned abundance outputs from a late-generation snapshot representing long-term demographic outcomes under regulation. For each replicate, abundances were summed across internal patches within each management area and size bin, separately for males and females. Replicate-level values were averaged to obtain mean size distributions. Male abundances were treated as negative values and female abundances as positive values to visualize population pyramids. Individuals below the legal size limit (25 cm) were distinguished from those at or above the limit.

#### Natal origin composition

Larval connectivity was assessed using natal origin data. For each current patch, individuals were grouped by natal patch of origin and summed across replicates. Counts were normalized within each current patch to obtain proportional contributions of natal origins. Natal patches outside the internal domain were pooled into a single category representing external sources.

#### Sex-specific differences in abundance

Sex-specific differences in abundance were evaluated using replicate-level counts. Individuals were grouped into two size classes (<25 cm and 25–35 cm). Male and female abundances were summed within each replicate for each combination of dispersal regime, MPA layout, and area type. Differences were calculated as male minus female abundance (M − F). One-sided Wilcoxon signed-rank tests were used to test whether male abundance was lower than female abundance. P-values were adjusted using the Benjamini–Hochberg procedure, and only comparisons with a false discovery rate below 0.01 were considered.

## Results

### Temporal dynamics of population size

Across all dispersal scenarios and spatial management layouts, total population size declined following the onset of fishing at generation 50 and began to recover after the implementation of regulation at generation 100 (**Figure 2**). The magnitude and timing of recovery, however, differed substantially among dispersal regimes and MPA configurations. A linear mixed□effects model of final population size revealed a significant three□way interaction among dispersal regime, MPA layout, and area type (Type III ANOVA, p < 0.001), indicating that population recovery depended on the interaction between connectivity structure and spatial configuration.

**Figure 2.**
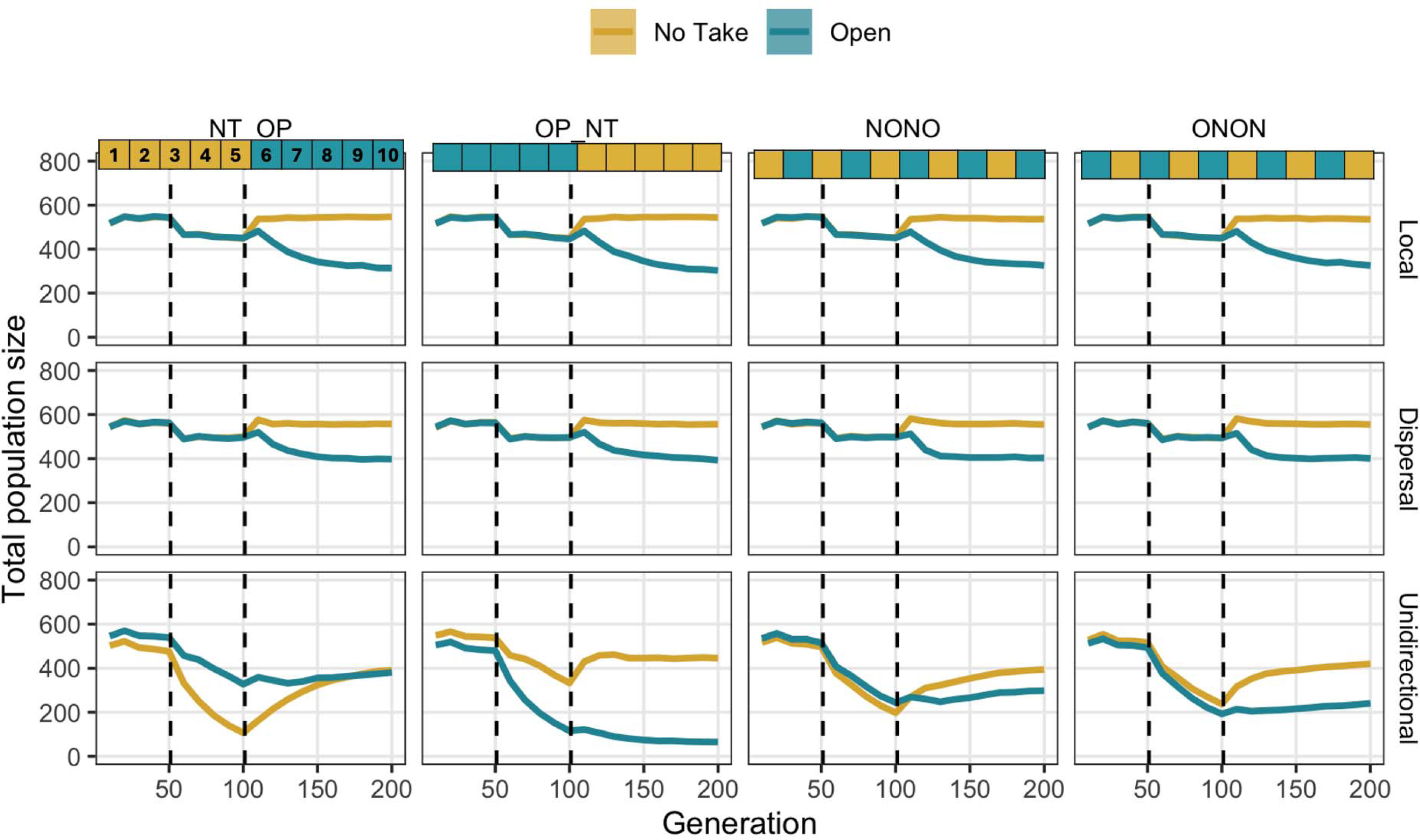
Population dynamics under alternative dispersal scenarios and spatial management layouts. Total population size through time, aggregated across internal patches and averaged across replicates, is shown for open (teal) and no-take (gold) areas. Columns correspond to four spatial management layouts (**NT_OP**, **OP_NT**, **NONO**, **ONON**), differing in the spatial arrangement of open and no-take patches along the coast. Rows correspond to three larval dispersal scenarios (**Local**, **Dispersal**, **Unidirectional**). Solid lines indicate mean population size across replicates, and shaded ribbons represent ±1 standard error. Vertical dashed lines indicate the onset of fishing (generation 50) and the onset of regulation (generation 101). Prior to regulation, all patches are subject to unrestricted harvest. After regulation begins, no-take patches are fully closed to fishing, whereas open patches allow harvest only within the legal size slot (25–32 cm) and exclude egg-bearing females from harvest.

Overall population recovery was stronger under the Local and Dispersal scenarios than under the Unidirectional dispersal scenario. No-take areas generally supported higher population sizes than open areas throughout the simulation, with one notable exception: under unidirectional dispersal, no-take patches placed upstream (NT_OP) did not consistently outperform open patches.

Under symmetric or weakly directional dispersal (Local and Dispersal scenarios), differences among MPA layouts were relatively minor. Population trajectories in open and no-take areas followed broadly similar patterns across layouts, indicating that spatial arrangement of protection had limited influence when larval exchange was local or approximately bidirectional.

In contrast, under unidirectional larval dispersal, population trajectories diverged markedly among spatial management layouts. In the NT_OP configuration, the no-take area experienced a pronounced decline immediately after fishing began, reaching a minimum around the time regulation was introduced, followed by a gradual and sustained recovery. Over the same period, the open area declined more slowly and remained at an intermediate population size throughout the post-regulation phase. This asymmetric response reflects strong downstream coupling, whereby upstream patches exported larvae downstream but received little replenishment in return.

In the OP_NT configuration, population recovery was substantially delayed. Open areas exhibited a sharp decline following the onset of fishing and showed little recovery even long after regulation was implemented. Although no-take areas increased in abundance after regulation, this increase did not translate into rapid recovery of adjacent open areas, resulting in a persistent disparity between management types.

The alternating layouts (NONO and ONON) showed intermediate dynamics relative to NT_OP and OP_NT. In both cases, populations in open and no-take areas declined strongly after fishing began and recovered partially following regulation. Recovery trajectories were slower than in NT_OP but more pronounced than in OP_NT, with neither area consistently dominating in post-regulation population size.

### Patch-level population size at equilibrium

After the regulation, patch-level population sizes differed systematically between dispersal regimes and management types (**Figure 3**). The qualitative patterns described below were consistent across generations, from generation 110 to generation 200, and generation 140 is shown as a representative example.

**Figure 3.**
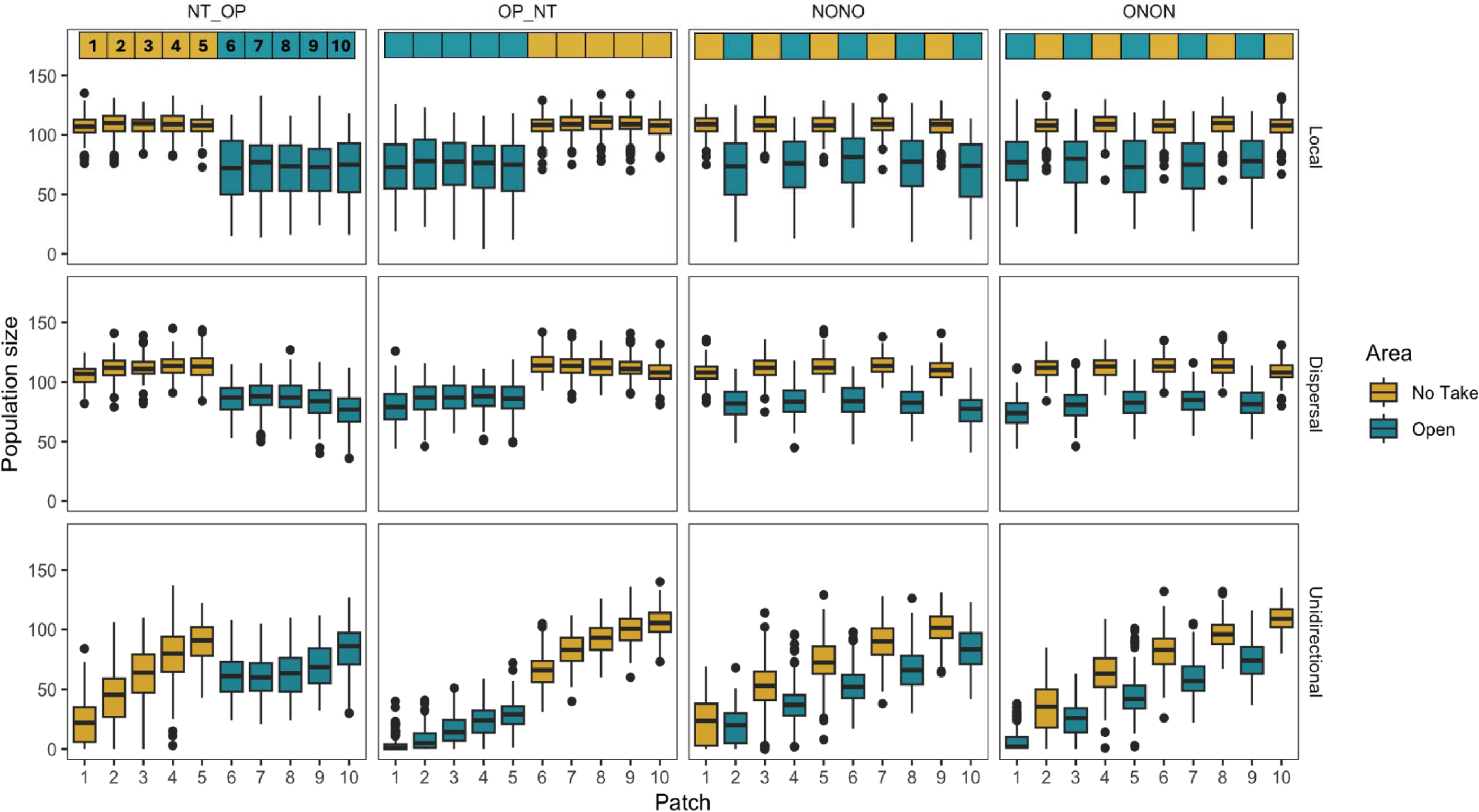
Patch-level population size distributions under dispersal scenarios and MPA layouts. Boxplots show the distribution of patch-level population size across replicate simulations for internal patches (1–10). Panels are arranged by larval dispersal scenario (rows: Local, Dispersal, Unidirectional) and spatial management layout (columns: NT_OP, OP_NT, NONO, ONON). Within each panel, boxplots are shown separately for no-take patches (gold) and open patches (teal), as defined by the corresponding MPA layout. Boxes indicate the interquartile range with the median line; whiskers extend to 1.5×IQR and points denote outliers.

Under the Local and Dispersal scenarios, population sizes in open areas were consistently lower than those in no-take areas across all spatial layouts and patches. This pattern was robust to spatial arrangement, indicating a persistent difference between management types when connectivity was local or bidirectional. Patch-level population size was significantly lower in open than in no-take areas under both Local and Dispersal scenarios (linear mixed-effects model, p < 0.001), and this difference was consistent across spatial layouts. Under unidirectional dispersal, population size increased with downstream position (p < 0.001), with a significant interaction between position and management type, indicating strong spatial structuring of abundance along the coast.

Under unidirectional dispersal, spatial position along the coastline strongly structured population sizes. When open patches were located downstream, open-area populations showed partial recovery, with higher abundances observed in downstream patches. Conversely, when open patches were located upstream, open-area populations remained low across patches and showed little evidence of recovery. Across all unidirectional layouts, population size increased with downstream position, with downstream patches supporting larger populations than upstream patches regardless of management type.

### Size structure at the final generation

At generation 200, no-take areas consistently supported higher abundances of large individuals (≥25 cm) than open areas across all spatial management layouts **(Figure 4)**. In contrast, open areas were dominated by smaller individuals (<25 cm), indicating persistent truncation of size structure relative to no-take zones. Differences in size structure between no-take and open areas were particularly pronounced under the NT_OP and OP_NT layouts, where contrasts in the abundance of large individuals were strongest. Under these configurations, open areas showed limited representation of harvestable size classes compared to adjacent no-take areas. Consistent with these patterns, statistical analyses at generation 200 revealed sex-specific differences in the abundance of large individuals. In a subset of dispersal scenario × layout × area-type combinations, the abundance of individuals in the 25–35 cm size class was significantly lower in males than in females after correction for multiple testing (one-sided Wilcoxon signed-rank tests, FDR < 0.01; **Table 1**, **Figure 4**).

**Figure 4.**
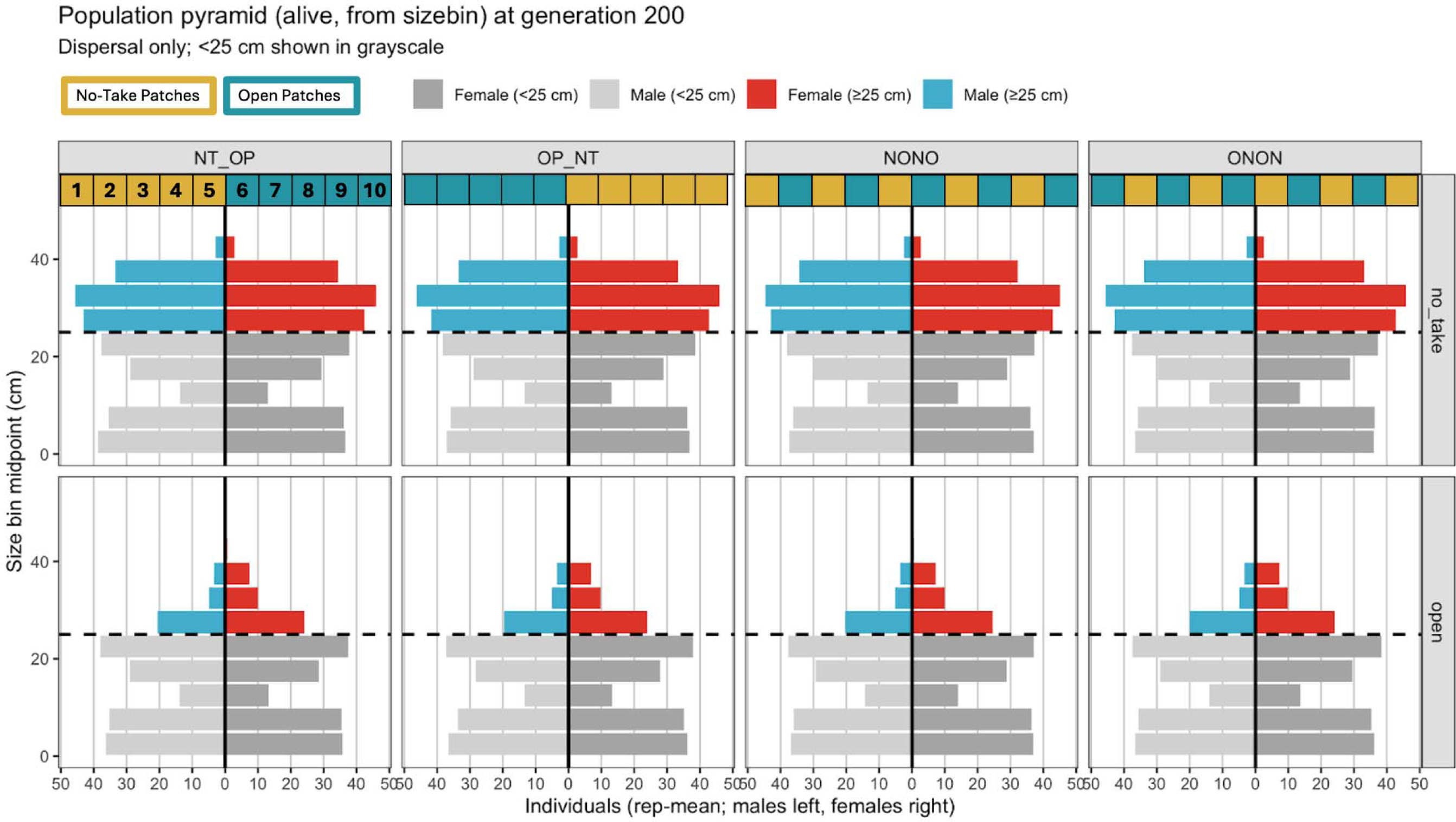
Size-based population structure (population pyramid) at generation 200 under the Dispersal scenario. Each panel shows the average number of live individuals per size class (5-cm bins) by sex and management area (no-take vs open), for each spatial layout. Individuals below the legal size limit (25 cm; shown in grayscale) are distinguished from those at or above the limit (colored: red for females, blue for males.

**Table 1.**
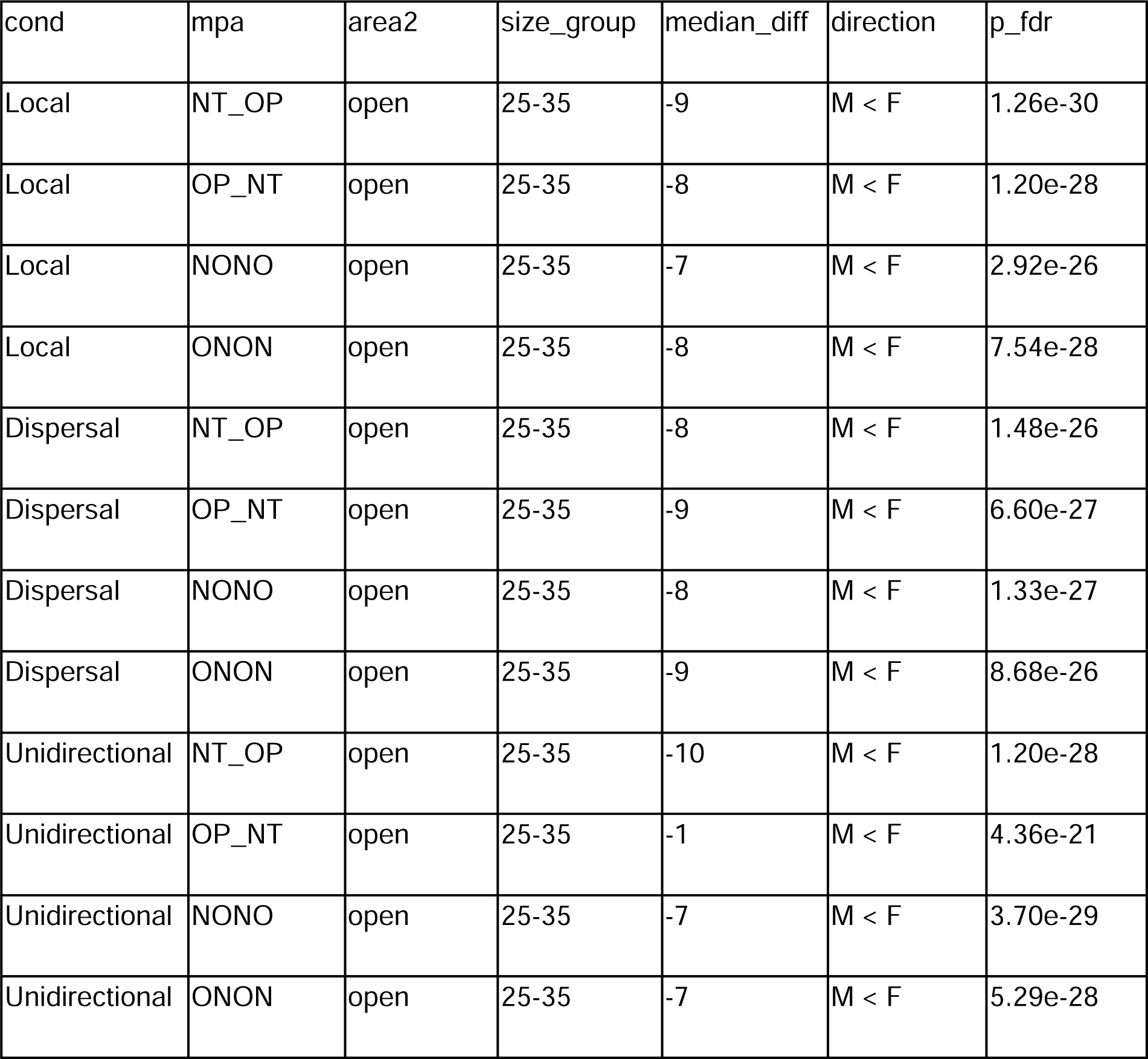
Sex-specific differences in abundance at generation 200. Sex-specific differences in abundance at generation 200 across dispersal scenarios (cond), spatial management layouts (mpa), and area types (area2). For each combination, replicate-level counts were summed within two size classes (<25 cm and 25–35 cm) and compared between males and females using one-sided Wilcoxon signed-rank tests (alternative hypothesis: M < F). P-values were adjusted for multiple testing using the Benjamini–Hochberg procedure, and only comparisons with FDR < 0.01 are reported. Median differences are shown as M − F.

### Natal origin and connectivity patterns

Across all MPA configurations, the natal origin composition of individuals within each patch closely reflected the underlying dispersal regime (**Figure 5**). Clear contrasts were observed among the Local, Dispersal, and Unidirectional scenarios.

**Figure 5.**
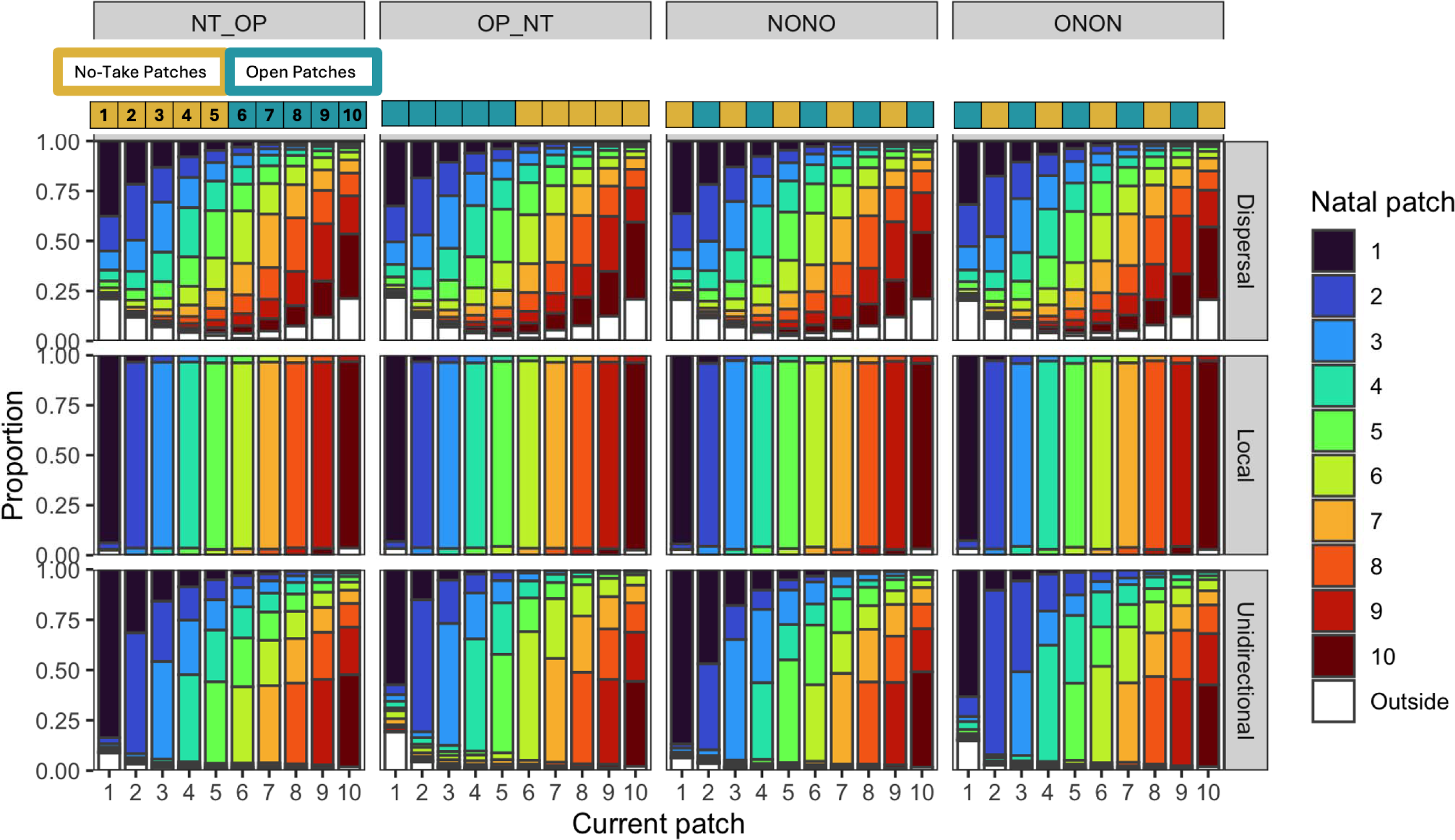
Natal origin composition of individuals across patches. Stacked bar plots show the proportional contribution of natal patches to the population present in each current patch (1–10) at generation 200. Colors indicate natal patch of origin; patches 0 and 11 are pooled and shown as “Outside”. Columns correspond to MPA configurations (NONO, ONON, OP_NT, NT_OP), and rows represent dispersal regimes (Dispersal, Local, Unidirectional). Bars are aggregated across replicates and normalized within each current patch to sum to one.

Under the Local dispersal regime, individuals in each patch were almost exclusively of local natal origin, indicating near-complete self-recruitment regardless of MPA configuration. In the Dispersal regime, patches contained a mixture of natal origins, with contributions from multiple neighboring patches, resulting in smooth spatial gradients in natal composition.

In contrast, the Unidirectional regime generated strongly asymmetric natal origin patterns. Downstream patches were dominated by individuals originating from upstream patches, consistent with directional larval connectivity and highlighting the strong spatial coupling imposed by unidirectional dispersal.

## Discussion

In this study, we used a spatially explicit individual-based model to clarify how larval dispersal structure and the spatial placement of no-take zones jointly determine the outcomes of marine protection in a sedentary coastal species. We show, first, that no-take zones consistently increase population abundance, promote larger size structure, and enhance reproductive potential relative to fished areas, confirming their role as demographic and reproductive refuges. Second, we demonstrate that the magnitude and spatial expression of these benefits depend critically on larval dispersal directionality. When dispersal is local or approximately symmetric, population responses are largely insensitive to the spatial configuration of protection. In contrast, under strongly unidirectional dispersal, management outcomes depend strongly on reserve placement, with pronounced differences among alternative spatial layouts. Together, these results demonstrate that the effectiveness of spatial protection cannot be evaluated independently of connectivity structure when dealing with species with pelagic larvae, and that alignment between larval dispersal direction and reserve placement is a key determinant of MPA performance in linear coastal systems.

Under unidirectional larval dispersal, the effectiveness of no-take zones emerges from a clear mechanistic asymmetry in larval supply and retention. Because larval export is strongly biased downstream, upstream no-take zones primarily function as sources of larvae that are transported away from the protected area. As a consequence, despite reduced fishing mortality on adults, upstream reserves experience limited self-recruitment and therefore fail to sustain high local population sizes following exploitation. In contrast, downstream no-take zones benefit simultaneously from larval import originating upstream and from local retention within the protected area. This dual advantage allows downstream reserves to accumulate recruits, rebuild abundance, and maintain a larger size structure. The resulting asymmetry in natal origin composition, with downstream patches dominated by upstream-derived larvae, directly links dispersal directionality to spatial differences in demographic recovery. Thus, under unidirectional connectivity, reserve placement determines whether protection intercepts larval supply or merely exports reproductive output, explaining why spatial configuration becomes a critical determinant of management success. These mechanisms are consistent with previous theoretical and empirical work showing that spatial configuration relative to connectivity strongly conditions MPA performance, particularly in systems with asymmetric dispersal (Costello *et al*., 2010; Ross *et al*., 2017).

Previous work has established that no-take MPAs frequently generate positive local responses, while larval dispersal plays a central role in determining whether these benefits extend beyond reserve boundaries. Biophysical and oceanographic modelling studies have demonstrated that larval connectivity is often highly asymmetric, with persistent source–sink dynamics driven by prevailing currents, bathymetry, and larval behaviour (Beng *et al*., 2025; Saito *et al*., 2025). These studies provide critical, system-specific descriptions of realised dispersal pathways and identify reserves that act as larval sources, sinks, or stepping stones within broader networks. In parallel, theoretical and applied work has highlighted that spatial information and connectivity patterns can fundamentally alter the performance of spatial management strategies (Costello *et al*., 2010; Balbar and Metaxas, 2019; Ovando *et al*., 2021; Nuno *et al*., 2024; Preston *et al*., 2025). However, most existing approaches either focus on describing connectivity patterns using high-resolution hydrodynamics or infer dispersal outcomes indirectly from genetic or demographic data. By contrast, the present study adopts a deliberately simplified, life-history–explicit modelling framework to disentangle these effects mechanistically. Rather than reproducing site-specific circulation, we use a minimal one-dimensional patch model to clarify how directional larval flow alters the balance between self-recruitment and export under alternative reserve layouts.

The biological relevance of these findings is particularly relevant to European lobster and other sedentary coastal invertebrates with planktonic larvae (Mare, 1942; Murray, 2007; Brummer and Kučera, 2022). European lobster exhibits strong adult site fidelity, such that direct spillover of adults from protected areas is limited, while population replenishment depends primarily on the production, dispersal, and settlement of larvae (Ellis *et al*., 2015b). In this context, size-dependent fecundity plays a critical role, as larger females are thought to contribute disproportionately to reproductive output, and protection from fishing allows individuals to survive to sizes with higher egg production (Flores *et al*., 2021; Coyle and Gnanalingam, 2026). Regulations that protect egg-bearing females further reinforce this effect by safeguarding reproductive output within protected areas (Sørdalen *et al*., 2022). However, our results suggest that adult protection alone is insufficient to ensure population recovery beyond reserve boundaries when larval dispersal is directional. Instead, effective spatial protection requires reserve placement that promotes larval retention or intercepts larval supply, thereby linking reproductive output to local or downstream recruitment. This combination of sedentary adult behaviour, size-dependent reproduction, and pelagic larval dispersal is common to many decapod crustaceans and other benthic invertebrates (Jablonski and Lutz, 1983; Strathmann, 1990), suggesting that the mechanisms identified here are likely to generalise beyond European lobster to a broader class of species with similar life-history strategies.

From a management perspective, our results highlight that the effectiveness of no-take zones cannot be evaluated solely on the basis of their size or level of protection, but must be considered in relation to patterns of larval connectivity. In systems where dispersal is directional, reserve placement relative to dominant larval flow is likely to be a key determinant of whether protected areas function as sources of recruitment or primarily export reproductive output. Incorporating even coarse information on dispersal direction into spatial planning may therefore improve the performance of MPA networks, particularly in linear coastal systems such as fjords, archipelagos, and narrow continental shelves.

Future work could integrate empirical estimates of lobster connectivity from genetic data or biophysical dispersal models to evaluate how the mechanisms identified here play out under realistic circulation patterns. Adult movement in European lobster is known to be highly limited and can be directly quantified through trap-based observations and tagging studies, providing relatively robust information on post-settlement connectivity (Øresland and Ulmestrand, 2013). In contrast, larval dispersal occurs during a planktonic phase that cannot be directly observed, and remains poorly quantified in many management contexts(Schmalenbach and Buchholz, 2010). Genetic approaches (Morales-González *et al*., 2019; van der Reis *et al*., 2022) could therefore provide critical insight into realised larval connectivity and source–sink dynamics, allowing clearer evaluation of whether existing or proposed MPAs are positioned to retain or intercept larval supply and to support population replenishment beyond reserve boundaries.

## Author Contribution

MS designed the study, developed and implemented the individual-based model, performed all simulations and analyses, and led the writing of the manuscript. LC and TH contributed domain expertise on European lobster biology, life-history traits, and fisheries management context, and provided conceptual input on the ecological and management questions addressed. All authors contributed to the manuscript revision.

## Data Availability Statement

All simulation outputs generated during this study are available in a public repository. Source code and analysis scripts are accessible at: https://github.com/mariesaitou/paper_2025-/tree/main/lob_SLiM

## Acknowledgments

Computational resources were provided by the Orion high-performance computing cluster at the Norwegian University of Life Sciences.

We acknowledge the NMBU lobster monitoring program for providing background material that helped frame the context of this study. We want to thank Stein R. Moe, Jonathan Colman, Linda Eikaas, Nora Colman, Odd Arne Sørensen and Knut Hove for their implications in the NMBU lobster monitoring program.

